# Instructor-learner brain coupling discriminates between instructional approaches and predicts learning

**DOI:** 10.1101/704239

**Authors:** Yafeng Pan, Suzanne Dikker, Pavel Goldstein, Yi Zhu, Cuirong Yang, Yi Hu

## Abstract

The neural mechanisms that support naturalistic learning via effective pedagogical approaches remain elusive. Here we use functional near-infrared spectroscopy to measure brain activity from instructor-learner dyads simultaneously during dynamic conceptual learning. We report that brain-to-brain coupling is correlated with learning outcomes, and, crucially, appears to be driven by specific scaffolding behaviors on the part of the instructors (e.g., asking guiding questions or providing hints). Brain-to-brain coupling enhancement is absent when instructors use an explanation approach (e.g., providing definitions or clarifications). Finally, we find that machine-learning techniques are more successful when decoding instructional approaches (scaffolding vs. explanation) from brain-to-brain coupling data than when using a single-brain method. These findings suggest that brain-to-brain coupling as a pedagogically relevant measure tracks the naturalistic instructional process during instructor-learner interaction throughout constructive engagement, but not information clarification.

## 1. Introduction

Humans have evolved the ability to learn through social interaction with others (e.g., an instructor), an important skill that serves us throughout our lifespan (Verga and Kotz, 2019; Pan et al., 2018). Such interactive learning is thought to be facilitated by instructional tools (Driscoll and Driscoll, 2005), like demonstrating rules or providing examples for practice. Verbal instruction has been shown to play an enabling and modulatory role in learning at multiple levels, ranging from functional brain re-organization (e.g., Hartstra et al., 2011; Olsson and Phelps, 2007; Ruge and Wolfensteller, 2009) to learning performance optimization (e.g., Clark and Mayer, 2016; Wolfson et al., 2014). However, despite the dynamic and interactive nature of instruction-based learning, neurobiological research investigating learning through instruction has been mostly limited to controlled laboratory studies – stripped from any real-time interaction between the learner and the instructor (e.g., Ruge and Wolfensteller, 2009) – and have often ignored the role of different instruction approaches (e.g., Holper et al., 2013). As a result, the brain mechanisms that support dynamic interactive learning remain understudied, and thus poorly understood.

Recent methodological advances (Brockington et al., 2018; for a review, see Hasson et al., 2012) have allowed researchers to begin investigating the neural basis of naturalistic instruction-based learning (Bevilacqua et al., 2019; Dikker et al., 2017; Liu et al., 2019; Pan et al., 2018). These studies have suggested that the interaction between instructor and learner is reflected in the extent to which brain activity becomes ‘coupled’ between them (Bevilacqua et al., 2019; Holper et al., 2013; Pan et al., 2018; Zheng et al., 2018). For example, brain-to-brain coupling has been reported to reliably predict the success of social interactive learning (Pan et al., 2018). However, while some studies have shown such a relationship between brain-to-brain coupling and learning outcomes (e.g., Holper et al., 2013; Liu et al., 2019; Pan et al., 2018; Zheng et al., 2018), others did not in fact observe a correlation between teacher-student brain-to-brain coupling and content retention (e.g., Bevilacqua et al., 2019). One potential limitation of most prior studies on learning concerns that they only focused on the average brain-to-brain coupling across the entire teaching session and its relation with learning outcomes (Davidesco et al., 2019). It is possible that linking specific moments of brain-to-brain coupling (such as those associated with certain instructional behavior) to learning might yield complementary useful information (Pan et al., 2018).

Here, we further investigated the functional significance of brain-to-brain coupling in learning and instruction. In addition to examining whether brain-to-brain coupling between instructors and learners can predict learning outcomes, we asked whether brain-to-brain coupling can be used to classify instructional dynamics during interactive learning. Such a finding would suggest that brain-to-brain coupling may be a pedagogically informative implicit measure that tracks learning throughout ongoing dynamic instructor-learner interactions.

We distinguished two instructional strategies (explanation vs. scaffolding), derived from two distinct pedagogical approaches to the role of instruction in instructor-learning interactions. First, the “explanation-based” approach assumes that learning emerges as a result of information clarification, which serves to enhance learners’ comprehension (Chi, 2013; Duffy et al., 1986). In this framework, instructional modulation of learning is driven by meaningful explanatory information. A second line of instructional approaches emphasizes the importance of supportive scaffoldings provided by the instructor. Scaffolding behaviors include asking key questions (e.g., asking learners their understanding of a core concept) and providing hints (e.g., giving an analogy of the learning content) that are aimed at redirecting learners’ actions and understanding (Van de Pol et al., 2010). Scaffolding foregrounds bidirectional communication and information sharing – both instructors and learners are involved in a two-way dynamic process of receiving and sending out information.

In addition to instructional strategy, adaptive behavior on the part of the instructor has also been shown critical for interactive learning (Chi, 2013; Chi and Roy, 2010). That is, the instructor provides personalized guidance based on the learner’s current level of knowledge (Wass and Golding, 2014). We therefore added a second dimension to our study design where half of the instructors were informed of the learner’s knowledge level based on their performance on a pre-test (personalized instruction) and half of them were not informed (non-personalized instruction).

Twenty-four instructor-learner dyads participated in a concept learning task, during which their brain activity was recorded simultaneously with functional near-infrared spectroscopy (fNIRS; Cheng et al., 2015; Pan et al., 2017; Zheng et al., 2018). Brain-to-brain coupling between instructors and learners was first estimated using Wavelet Transform Coherence (Grinsted et al., 2004), and then correlated with learning outcomes. A video coding analysis allowed us to parse whether the brain-to-brain coupling in instructor-learner dyads was specifically driven by certain instructional behavior. Finally, to identify to what extent scaffolding strategies can be distinguished from explanation strategies in the neural data, we used a decoding analysis. We employed the same decoding approach on both brain-to-brain coupling and individual brain data to explore the possible added value of a two-brain vs. single-brain analysis.

## 2. Methods

### 2.1. Participants

Twenty-four dyads (n = 48, all females, mean age = 21.46 ± 2.75 years) were recruited to participate in the study. Each dyad consisted of one learner and one instructor. Each instructor taught the learner in a one-to-one way. The instructors (mean age = 22.58 ± 2.75 years) had all received graduate training in psychology, had at least 1-year of instructional experience, and were familiar with the learning content, whereas the learners (mean age = 20.33 ± 2.30 years) in our sample majored in non-psychology related fields and had not been exposed to the content. All participants were healthy and right-handed and were recruited through advertisements. Each participant gave informed consent prior to the experiment and was paid for participation. The study was approved by the University Committee of Human Research Protection (HR 044-2017), East China Normal University.

### 2.2. Tasks and materials

The task used in the present fNIRS-based hyperscanning study was a conceptual learning task, which involved mastering two sets of materials, each explaining four psychological terms pertaining to an overarching concept. The material was chosen to be novel and attractive to non-psychology majors and teachable within 10 – 20 minutes. The sets centered around the concepts of *reinforcement* and *transfer.* These concepts were chosen from a classic national standard textbook (Educational Psychology: A Book for Teachers, Wu & Hu, 2003). These two concepts belong to the similar topic (i.e., learning psychology) and occupy a similar instructional period (i.e., 1~2 sessions). The *reinforcement* set consisted of teaching positive reinforcement, negative reinforcement, punishment, and retreat (Set 1), and *transfer* consisted of near-transfer, far-transfer, lateral-transfer, and vertical-transfer (Set 2). This design allowed us to provide different learning content for the two within-participant instructional strategies (i.e., scaffolding vs. explanation), without repeating any content. Learning outcomes did not differ between the two sets of concepts, and were thus pooled together in the results reported below.

All instructors were informed and trained by experimenters two days prior to the experiment. Training examples were selected from the textbook’s training section. Each example consisted of instructional goals, instructional difficulties, general instructional processes, and detailed instructional scripts. Such instructional scripts were composed and adapted with the help of two psychological experts with at least 20 years of instructional experience at the university level. Instructors were required to prepare instruction at home for 2 days. They then practiced with each other in the lab until they were satisfied with their own instructional performance in both the scaffolding and explanation conditions (they spent approximately the same amount of time training for both types of instructions). Then they demonstrated instruction to the experimenter in a one-to-one way until their performance met the established standard requirements: the length of teaching, the speed of speech, and consistency with the instructional processes and scripts (Liu et al., 2019).

### 2.3. Experimental factors

We manipulated one within-participant variable and one between-participant variable. The within-participant variable was the Instructional Strategy (scaffolding vs. explanation). Following the scripts, the instructor using a scaffolding strategy would guide the learner in a Q&A manner along the following lines (one representative example, translated from Chinese):

- *Instructor: How can one provide positive reinforcement?*
- *Learner: …… By rewarding positive behavior?*
- *Instructor: Bingo! Could you please give an example?*
- *Learner: My sister gave me some candies after I cleaned my room.*
……

For the explanation strategy, the instructor would explain each concept to the learner and provide examples. The following interaction provides a representative example of explanatory behavior:

- *Instructor: Positive reinforcement refers to rewarding goal-directed behavior to increase its frequency. Do you see what I mean?*
- *Learner: I am not sure whether I understand it correctly. Could you please explain it a bit more?*
- *Instructor: For example, my mom cooks my favorite food for me when I pass exams.*
- *Learner: That clarifies it.*
……

The between-participant variable was Instructional Personalization (personalized vs. non-personalized; i.e., whether the instructor customizes their instructions to the learner’s aptitude and ability as established via a pre-test). Instructions might be intrinsically personalized: for example, instructors often monitor learners’ comprehension and guide their understanding during face-to-face interactions. For instructors to be able to customize their instructions, learners have to inform them about their lack of understanding. Therefore, we exogenously manipulated

Instructional Personalization. For half of the participants (*n* = 12 dyads), the learner’s pre-test results (i.e., prior knowledge level) of the eight concepts (4 from Set 1 and 4 from Set 2) were provided to the instructor. The instructor was then asked to adapt their instruction to suit the needs of each learner (e.g., allocate more time to the teaching of a concept if the learner had difficulty learning it). For the non-personalized group (*n* = 12 dyads), the instructor was provided no information about the learner.

### 2.4. Procedures

The task included two blocks, each split into a resting-state phase and an interactive learning phase (**Fig. 1A**). The inter-block interval was approximately 1 minute. During the initial resting-state phase (3 min), both participants (sitting face-to-face, 0.8 meters apart) were asked to relax and to remain still. This 3-min resting phase served as the baseline.

**Figure 1.**
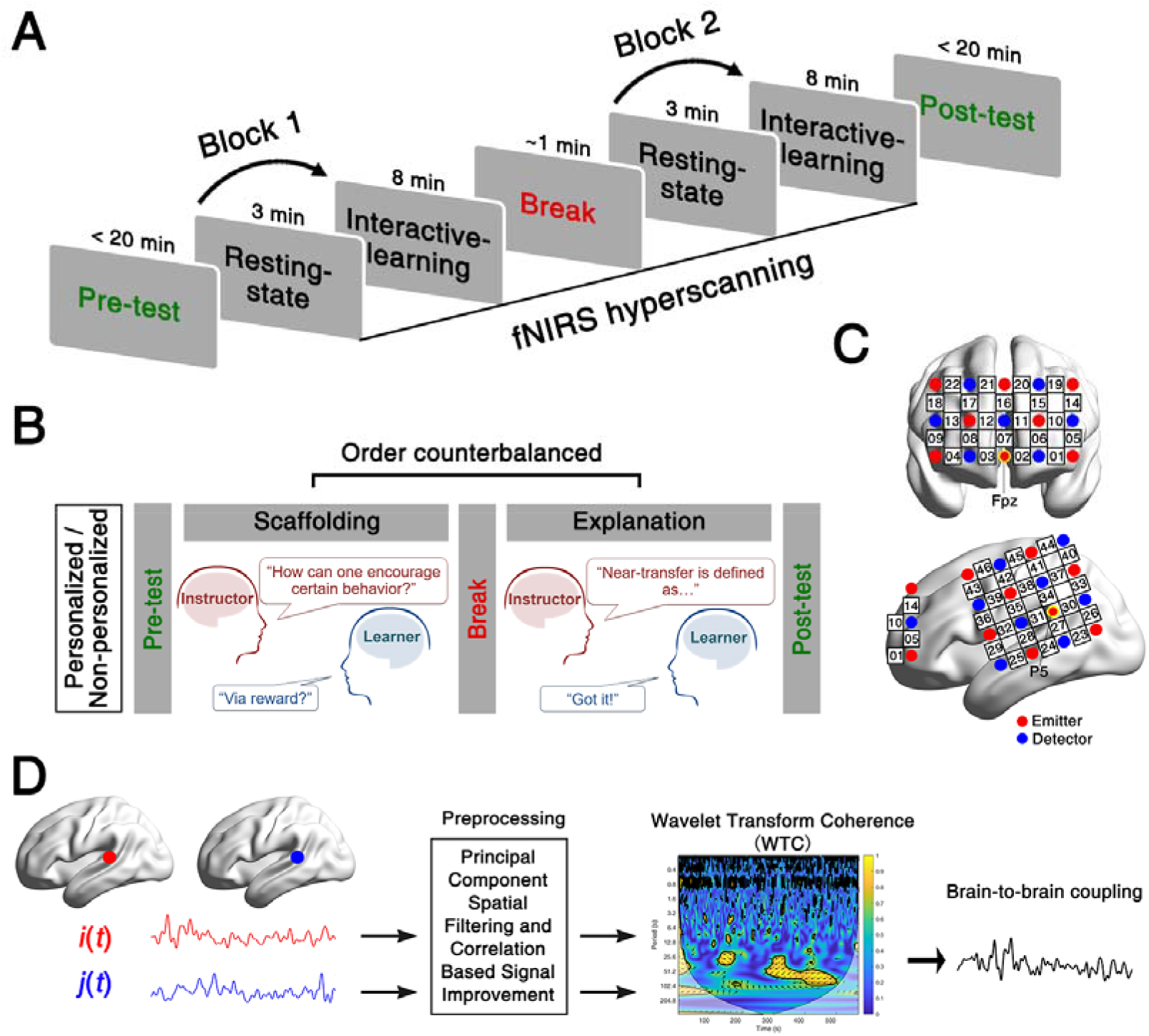
Experimental protocol, probe location, and brain-to-brain coupling analysis. **(A)** Experimental procedure. Before and after scanning, learners’ knowledge of the psychological concepts was evaluated. Brain activity from the instructor and the learner were acquired simultaneously using fNIRS, in two blocks, each starting with a 3-min rest (resting-state phase/baseline), followed by the instructor teaching concepts to the learner (interactive-learning phase/task). **(B)** Instructional Personalization and Instructional Strategies. Participants were randomly allocated to either personalized or non-personalized groups (Instructional Personalization). Within each instructor-learner dyad, scaffolding and explanation strategies were compared. **(C)** Optode probe set. The set was placed over prefrontal and left temporoparietal regions. **(D)** Overview of the brain-to-brain coupling analysis. Channel-wise raw time courses were extracted from both the instructor and the learner. After a battery of preprocessing, brain-to-brain coupling was estimated by Wavelet Transform Coherence between the two clean time courses. *i, j,* fNIRS signals of two participants of a dyad; *t*, time.

The resting-state phase was immediately followed by the interactive-learning phase (8 min), where the learner and instructor engaged in interactive learning either in a personalized (n =12 dyads) or non-personalized (n = 12 dyads) way (Instructional Personalization, **Fig. 1B**). For each group, the experimental procedure consisted of one of the following combinations of learning content and Instructional Strategy: (*i*) *reinforcement* with scaffolding (block 1) + *transfer* with explanation (block 2), (*ii*) *reinforcement* with explanation (block 1) + *transfer* with scaffolding (block 2). Block order was counterbalanced.

During the experiment, learners’ and instructors’ brain activity was recorded simultaneously via fNIRS-based hyperscanning at prefrontal and left temporoparietal regions (**Fig. 1C**). A digital video camera (Sony, HDR-XR100, Sony Corporation, Tokyo, Japan) was used to record the behavioral interactions between participant dyads. The acquisition of video data and fNIRS data was synchronized with a real-time audio-video cable connecting the camera to the ETG-7100 equipment. The camera recordings were used to classify (following the experiment) behavior as either scaffolding or explanatory behaviors.

### 2.5. Learning tests and outcome analysis

Learners’ knowledge of psychological concepts was tested immediately before the onset of the resting-state phase and after the end of the interactive-learning phase. Relevant to Reinforcement and Transfer, 8 definitions, 16 true-false items and 4 short answer questions were selected from textbooks to compose a testing bank. These items were randomly split into two halves, one for the pre-test and the other for the post-test. Results from 9 participants who were not involved in the fNIRS study showed that the difficulty levels did not differ between the pre- and post-tests (*t*_(8)_ = 0.01, *p* = 0.99). The learners had a time limitation of 20 min to finish each of the tests (Zheng et al., 2018).

The performance of learners in the pre- and post-tests was scored by two separate other raters who were blind to the group assignment. Three question types (i.e., definitions, true-false items, simple answer questions) were evaluated. For each learner, inter-coder reliability was calculated by the intra-class correlation on scores for definitions and simple answer questions (ranging from 0.77 to 0.91). Rating scores were averaged across the two raters. The sum of the judgments made on all three question types (for a given learner) was considered as the index of overall learning performance [maximum score: 4 (for 4 definitions) + 16 (for 8 true-false items) + 10 (for 2 simple answer questions) = 30 points). Pre-test scores did not differ between any of the conditions (*F*_s_ < 1.60, *p*s > 0.17). For all subsequent analyses, learning outcomes were quantified as the difference pre-learning scores and post-learning scores. A mixed-design repeated measures ANOVA was conducted on the learning outcomes, with Instructional Personalization (personalized vs. non-personalized) as a between-subject variable and Instructional Strategy (scaffolding vs. explanation) as a within-subject variable.

### 2.6. Image acquisition

An ETG-7100 optical topography system (Hitachi Medical Corporation, Japan) was used for brain data acquisition. The absorption of near-infrared light (two wavelengths: 695 and 830 nm) was measured with a sampling rate of 10 Hz. The oxyhemoglobin (HbO) and deoxyhemoglobin (HbR) were obtained through the modified Beer-Lambert law. We focused our analyses on the HbO concentration, for which the signal-to-noise ratio is better than HbR (Mahmoudzadeh et al., 2013). A number of fNIRS-based hyperscanning reports have used this indicator to compute of brain-to-brain coupling (e.g., Cheng et al., 2015; Dai et al., 2018; Jiang et al., 2012, 2015; Pan et al., 2017; Tang et al., 2015).

Two optode probe sets were used to cover each participant’s prefrontal and left temporoparietal regions (**Fig. 1C**), which have been previously associated with information exchanges between instructors and learners during interactive learning (Holper et al., 2013; Pan et al., 2018; Takeuchi et al., 2017; Zheng et al., 2018). One 3 × 5 optode probe set (eight emitters and seven detectors forming 22 measurement points with 3 cm optode separation) was placed over the prefrontal area. The middle optode of the lowest probe row of the patch was placed at Fpz (**Fig. 1C**), following the international 10-20 system (Okamoto et al., 2004). The middle probe set columns were placed along the sagittal reference curve. The other 4 × 4 probe set (eight emitters and eight detectors forming 24 measurement points with 3 cm optode separation) was placed over the left temporoparietal regions (reference optode was placed at P5, **Fig. 1C**). The correspondence between the NIRS channels (CHs) and the measured points on the cerebral cortex was determined using a virtual registration approach (Singh et al., 2005; Tsuzuki et al., 2007).

### 2.7. Imaging-data analyses

#### 2.7.1. Analysis step A: Brain-to-brain coupling

Data collected during the resting-state phase (3 min, served as the baseline) and the interactive-learning phase (8 min, served as the task) in each block were entered into the brain-to-brain coupling analysis (**Fig. 1D**). A principal component spatial filter algorithm was used to remove systemic components such as blood pressure, respiratory and blood flow variation from the fNIRS data (Zhang et al., 2016). To remove head motion artifacts, we used a “Correlation Based Signal Improvement” approach (Cui et al., 2010).

We then employed a wavelet transform coherence (WTC) analysis to estimate brain-to-brain coupling. The WTC of signals *i*(*t*) and *j*(*t*) was defined by:

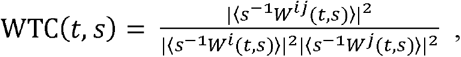

where *t* denotes the time, *s* indicates the wavelet scale, 〈·〉 represents a smoothing operation in time and scale, and *W* is the continuous wavelet transform (see Grinsted et al., 2004 for details). Our brain-to-brain coupling analysis was conducted in a data-driven manner and entailed three sub-steps:

##### Step 1: Does interactive learning lead to enhanced brain-to-brain coupling compared to baseline?

As a first step, we estimated whether brain-to-brain coupling was enhanced during the interactive learning task (estimated by WTC) compared to baseline. Time-averaged brain-to-brain coupling (also averaged across channels in each dyad) was compared between the resting phase (i.e. baseline session) and the interactive learning phase (i.e. task session) using a series of paired sample t-tests, one for each frequency band (frequency range: 0.01 – 1 Hz, Nozawa et al., 2016). This analysis yielded a series of *p*-values that were FDR corrected (*p* < 0.05). This analysis enables the identification of frequency characteristic, which help us determine the frequency of interest (FOI) for subsequent analyses.

To verify if the enhanced brain-to-brain coupling was dyad-specific, data from all 48 participants were reshuffled in a pseudo-random way so that 24 new dyads were created (e.g., time series from instructor #1 were paired with those from learner #3) (**Fig. 3E**). Then, the above brain-to-brain coupling analysis was performed again to obtain brain-to-brain coupling for pseudo-pairs.

**Figure 2.**
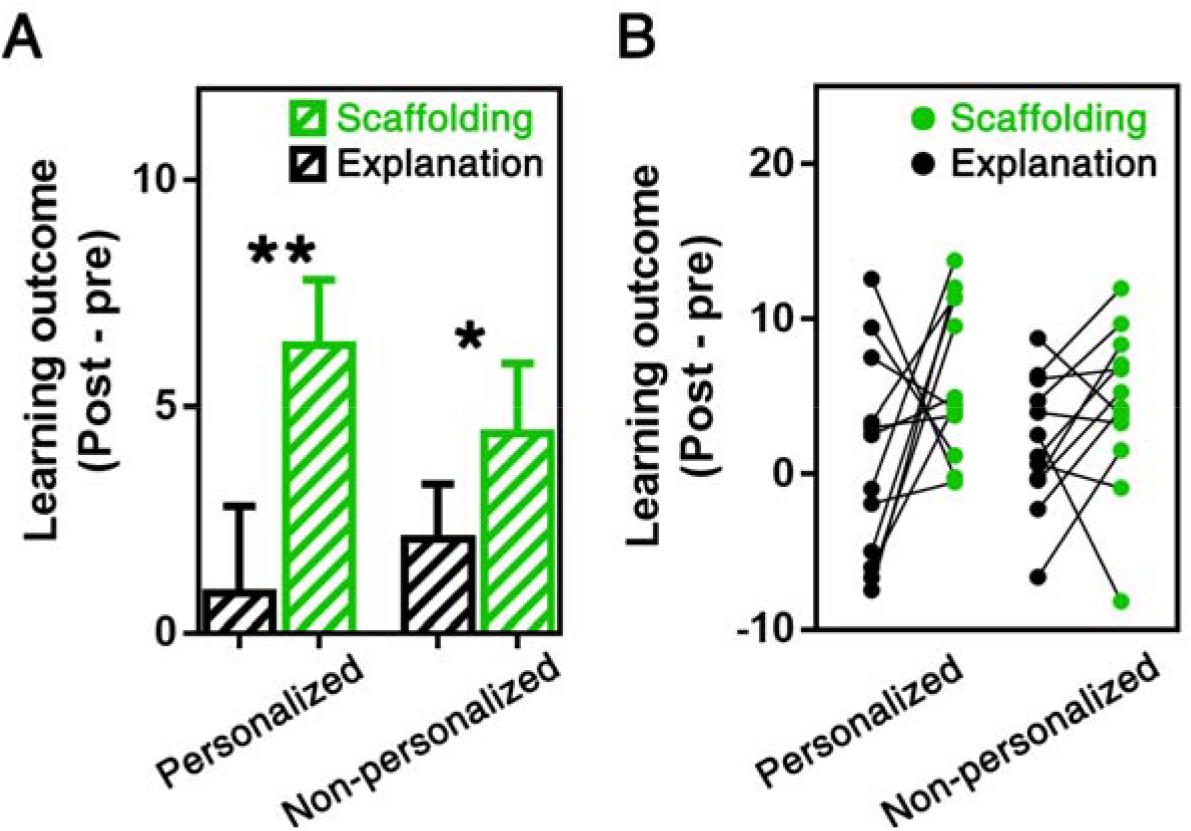
Learning outcomes in all conditions. **(A)** Group levels: in both personalized and non-personalized groups, learning outcomes for the scaffolding condition was significantly higher than the explanation condition. Learning outcomes are indexed by the change score (post-test score minus pre-test score). Error bars represent standard errors of the mean. **(B)** Corresponding graph for individual levels. \**p* < 0.05. \*\**p* < 0.01.

**Figure 3.**
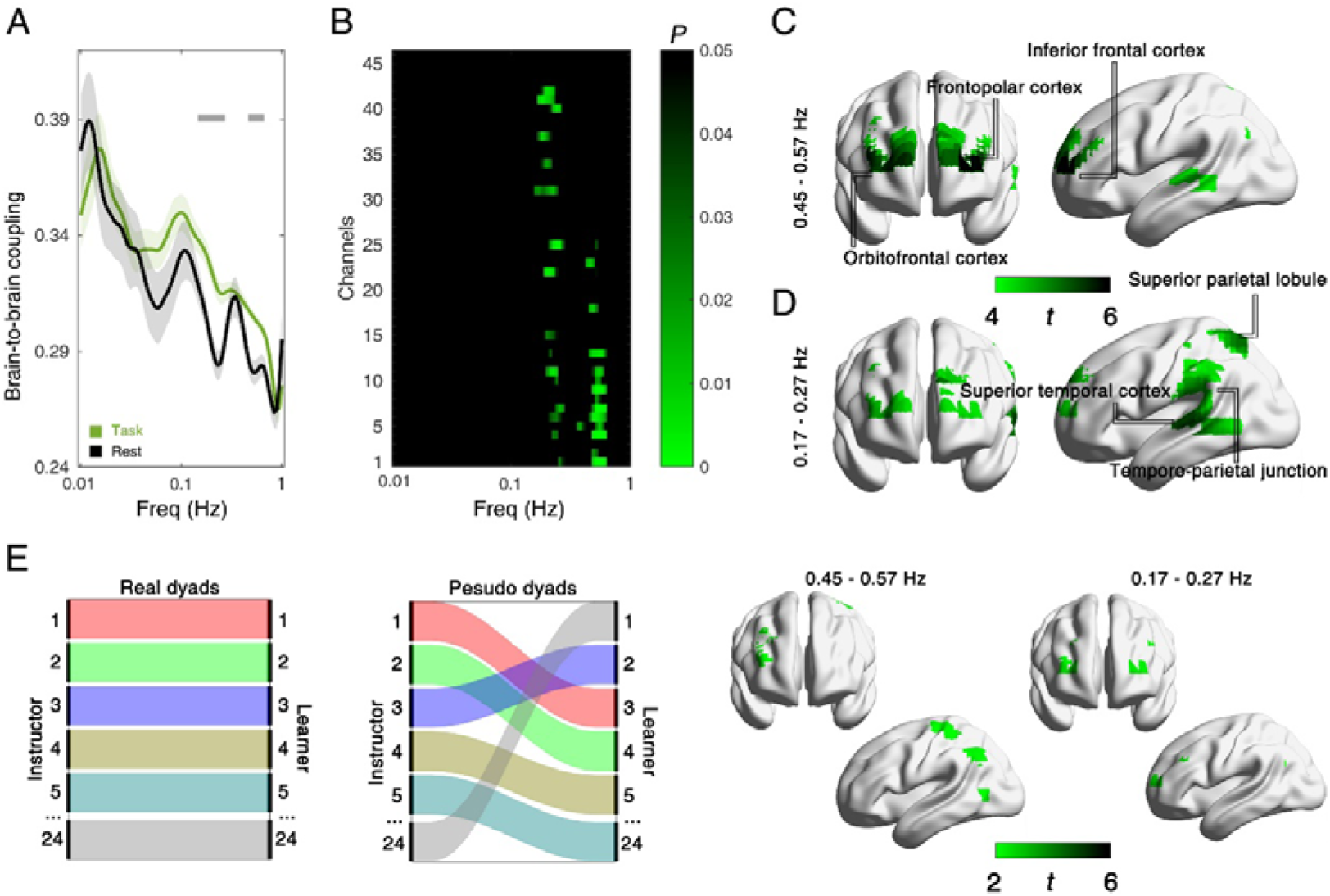
Interactive learning evokes frequency-specific widespread brain-to-brain coupling across all conditions. **(A)** Brain-to-brain coupling associated with the instruction session and the rest session for frequencies ranging between 0.01 and 1 Hz (all participants and channels’ data were averaged). Grey horizontal lines on the top indicate which frequencies show statistical differences (FDR controlled). **(B)** An FDR-corrected *P*-value map resulting from comparisons between instruction and rest (for each channel) across frequencies between 0.01 and 1 Hz. Interactive learning evokes frequency-specific widespread brain-to-brain coupling in all conditions across all dyads at 0.45 – 0.57 Hz **(C)** and 0.17 – 0.27 Hz **(D)**. **(E)** Control analyses confirmed that the enhanced brain-to-brain coupling shown in **(C)** and **(D)** was dyad-specific: no significant task-related coupling was detected in pseudo-dyads in either frequency band of interest (all real dyads were shuffled, resulting in 24 new pseudo dyads).

##### Step 2: Does task-related brain-to-brain coupling enhancement differ across the experimental conditions?

We averaged brain-to-brain coupling within each identified FOI and compared all conditions. We computed an index of task-related brain-to-brain coupling by subtracting the averaged coupling during the resting phase from that during the interactive learning phase. Fisher z transformation was applied to the task-related coupling values to generate a normal distribution. The resulting values for each channel were then submitted into an Instructional Strategy (scaffolding vs. explanation) × Instructional Personalization (personalized vs. non-personalized) mixed-design ANOVA. Parallel analyses were conducted separately in each FOI. The resulting *p* values were FDR-corrected for multiple comparisons. The results yielded *F* maps for each FOI. These *F* maps were visualized using BrainNet Viewer (Xia et al., 2013).

##### Step 3: Is condition-specific brain-to-brain coupling predictive of learning?

Finally, we assessed behavior-brain relationships. Pearson correlational analyses were employed to test the relationship between task-related brain-to-brain coupling from significant channels and learning outcomes.

#### 2.7.2. Analysis step B: Brain-to-brain coupling segmentation

Following the brain-to-brain coupling analyses, we grouped and averaged the adjacent CHs that showed significant brain-to-brain coupling as channels of interest. The time course of brain-to-brain coupling in the channels of interest was down-sampled to 1 Hz to obtain point-to-frame correspondence between the time series and video recordings (**Figs. 5A&B**).

**Figure 5.**
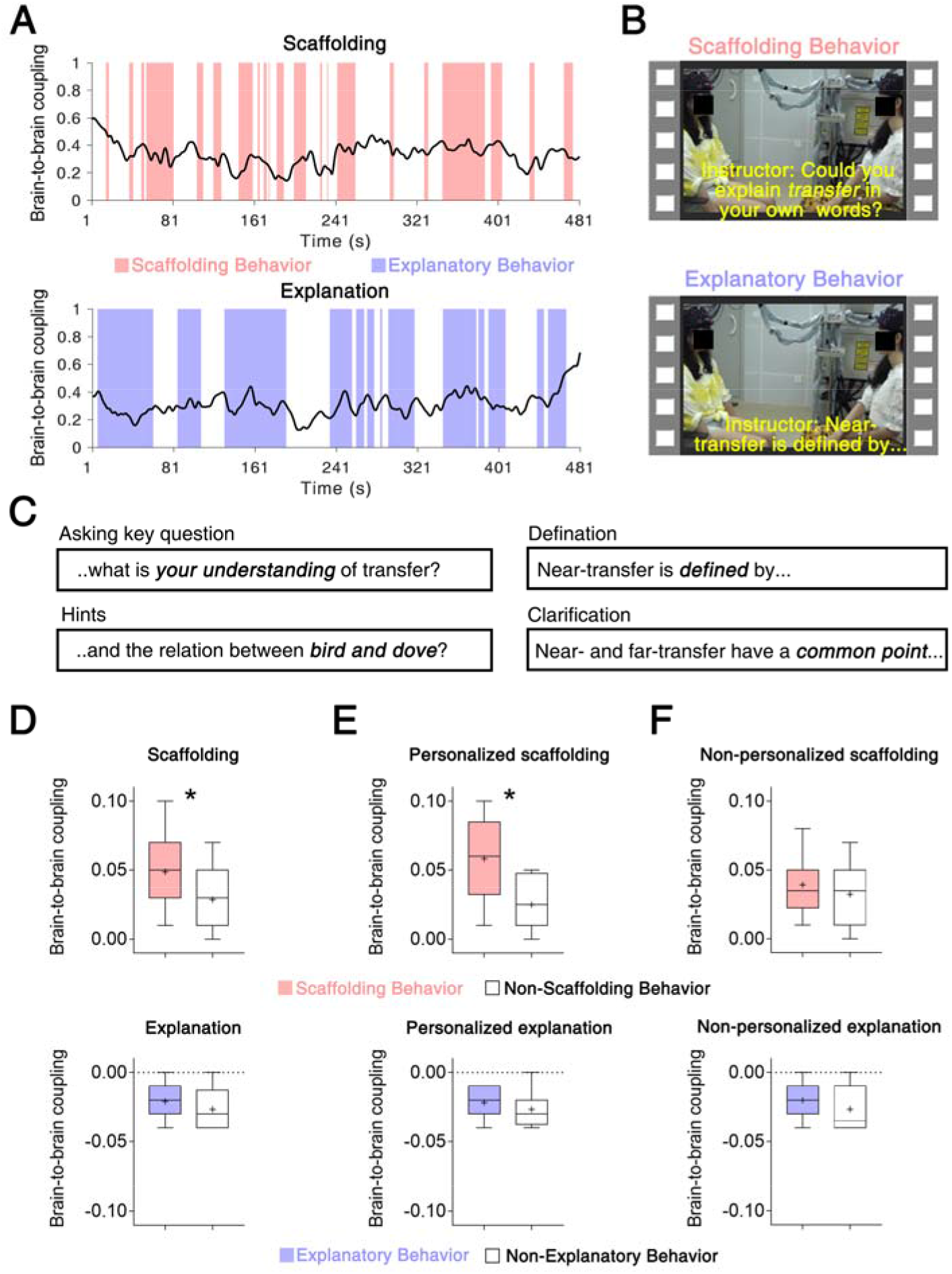
Video coding analysis reveals that brain-to-brain coupling is driven by specific instructional behaviors. **(A)** Time course of brain-to-brain coupling in the learning phase for one randomly selected dyad from the scaffolding and explanation conditions. Vertical panels denote the instructional behaviors: red panels indicate scaffolding behaviors; blue ones indicate explanatory behaviors. **(B)** Examples of each instructional behavior as coded from the video frames. **(C)** Example sentences from the video coding analysis for scaffolding behaviors (asking key questions and providing hints) and explanation behaviors (definition and clarification). Box plots of task-related brain-to-brain coupling (task minus rest) across the instructional behaviors in the scaffolding and explanation conditions **(D)**, in the personalized scaffolding and personalized explanation conditions **(E)**, and in the non-personalized scaffolding and non-personalized explanation conditions **(F)**. Crosses indicate the average brain-to-brain coupling across participant dyads. Error bars range from the min to the max value observed. \**p* < 0.05.

Two graduate students were recruited to independently code instructional behaviors in the interactive-learning phase using the video-recording data. The two coders underwent a weeklong training program by an educational expert (with 28 years of instructional experience in the field of education) to correctly identify instructional behaviors. Two types of instructional behaviors were categorized for each Instructional Strategy: for the scaffolding condition, there were (*i*) scaffolding behaviors, such as asking key questions, providing feedback and hints, prompting, simplifying problems, and (*ii*) other non-scaffolding instructional behaviors, i.e., those segments in the videos where scaffolding did not occur; for the explanation condition, there were (*i*) explanatory behaviors, such as giving detailed definitions, providing prefabricated materials, and information clarification, and (*ii*) other non-explanatory instructional behaviors, i.e., those segments in the videos where explanation did not occur.

Each one-second (s) video fragment (from the 8 minutes during the interactive-learning phase) was coded as either containing scaffolding behaviors or non-scaffolding instructional behaviors in the scaffolding condition; and as either consisting of explanatory behaviors or non-explanatory instructional behaviors in the explanation condition. For all coding activities, inter-coder reliability was calculated by the intra-class correlation (Werts et al., 1974). Inter-coder reliability was 0.87 for the scaffolding behaviors (vs. non-scaffolding instructional behaviors) in the scaffolding condition, and 0.81 for the explanatory behaviors (vs. non-explanatory instructional behaviors) in the explanation condition. If there was an inconsistency, the two coders discussed it and came to an agreement.

Based on the results of the coding procedures mentioned above, we categorized the segments of brain-to-brain coupling associated with different video-coded instructional behaviors (**Figs. 5A&B**). We subtracted brain-to-brain coupling during the rest session (baseline) from these segments of brain-to-brain coupling to obtain the task-related coupling. Contrasts between task-related brain-to-brain coupling associated with different video-coded instructional behaviors were obtained using a series of paired-sample *t*-tests.

#### 2.7.3. Analysis step C: Brain-to-brain coupling prediction

Finally, we explored whether brain-to-brain coupling allowed us to predict if an instructor employed the *scaffolding* or *explanation* strategy, using a decoding analysis (Dai et al., 2018; Jiang et al., 2015). The analysis details and strategies can be described as follows.

##### Classification features and labels

The time-averaged brain-to-brain coupling values at channels of interest were used as classification features. We first averaged the brain-to-brain coupling across the whole time series, resulting in time-averaged coupling for each channel. We focused on the channel(s) that exhibited significant task-related coupling (task vs. baseline; Goldstein et al., 2018). Instructional Strategies (i.e., *scaffolding* or *explanation)* were used as class labels.

##### Classification algorithm

Brain-to-brain coupling features were incorporated into a logistic regression algorithm. Logistic regression is a supervised machine-learning algorithm that has been previously used to predict behavioral measures with neuroimaging data (e.g., Ryali et al., 2010). The aim of logistic regression-based machine learning is to find the best fitting model that describes the relationship between the dichotomous features of the dependent variable and independent variables (Yan et al., 2004).

##### Classification performance

Classification performance was assessed using the standard metric of area under the receiver operating characteristic curve (AUC). The AUC is one of the most common quantitative indexes (Faraggi and Reiser, 2002; Hanley and McNeil, 1982), which illustrates the sensitivity and specificity for the classifier output. It has been successfully used to quantify the accuracy of the prediction in many neuroimaging studies (e.g., Cohen et al., 2018; Ki et al., 2016).

A permutation test was used to determine whether the obtained AUC was significantly larger than that generated by chance. Chance level of the AUC was determined by randomly shuffling the labels (*scaffolding* or *explanation)* for the brain-to-brain coupling values. Significant levels (*p* < 0.05) were calculated by comparing the correct AUC from the real labels with 10000 renditions of randomized labels.

##### Additional analyses

Finally, we tested whether decoding based on brain-to-brain coupling generated a better classification of instructional behavior than decoding based on individual brain activation. The raw fNIRS data were first preprocessed following the same procedure described in *Analysis Step A*. Clean (task-related) signals were then converted into *z*-scores using the mean and the standard deviation of the signals recorded during rest (baseline). Normalized intra-brain activity values at channels of interest in both instructors and learners were extracted as classification features. The parallel decoding analyses were then repeated as described above.

## 3. Results

### 3.1. Behavioral performance

A repeated measures ANOVA on learning outcomes with Instructional Strategy (Scaffolding vs. Explanation) as a within-dyad factor and Instructional Personalization (Personalized vs. Non-personalized) as a between-dyad factor revealed a main effect of Instructional Strategy (*F*_(1, 24)_ = 5.10, *p* = 0.03, *η*_partial_^2^ = 0.19), with the scaffolding strategy showing better learning outcomes than the explanation strategy (**Fig. 2**). There was no effect of Instructional Personalization on learning (*F*_(1, 24)_ = 0.82, *p* = 0.38) and there was no interaction between Instructional Personalization and Instructional Strategy (*F*_(1, 24)_ = 0.07, *p* = 0.79). In sum, learners who were taught using scaffolding retained more content from the instruction than learners who were taught using an explanation-based instructional strategy.

### 3.2. Brain imaging results

#### 3.2.1. Interactive learning induces frequency-specific widespread brain-to-brain coupling

In a first-pass data-driven analysis, we calculated brain-to-brain coupling in all conditions across the whole sample of 24 participant dyads to test whether interactive learning (i.e., task) was associated with enhanced brain-to-brain coupling compared to the resting-state session (i.e., baseline).

In terms of frequency characteristics, brain-to-brain coupling was significantly higher during the interactive learning phase than during rest for frequencies ranging between 0.45 – 0.57 Hz and 0.17 – 0.27 Hz (all FDR-corrected, **Fig. 3**). These two ranges were then chosen as frequencies of interest (FOIs) for subsequent analyses. These FOIs are out of the range of physiological responses associated with cardiac pulsation activity (~ 0.8 – 2.5 Hz) and spontaneous blood flow oscillations (i.e., Mayer waves, ~ 0.1 Hz).

Regarding spatial characteristics, task-related coupling enhancement was highest in the orbitofrontal cortex, frontopolar cortex, and inferior frontal cortex at 0.45 – 0.57 Hz (**Fig. 3C**), and along superior temporal cortex, temporoparietal junction, and superior parietal lobule at 0.17 – 0.27 Hz (**Fig. 3D**). We also observed widespread brain-to-brain coupling in adjacent regions, including prefrontal, temporal, and parietal areas. These results replicate previous research showing that social interactive learning (through instruction) induces brain-to-brain coupling in high-order brain regions (Holper et al., 2013; Pan et al., 2018; Zheng et al., 2018).

A control analysis confirmed that the patterns of brain-to-brain coupling (higher coupling associated with interactive learning compared to rest) were specific to the interaction between real instructor-learner dyads: pseudo dyads did not show higher brain-to-brain coupling during learning than rest (*p*s > 0.05, FDR controlled, **Fig. 3E**). Together, our first-pass results suggest that social interactive learning induces widespread brain-to-brain coupling. This coupling is concentrated in specific frequencies and only emerges in ‘real’ dyads (who are actually interacting).

#### 3.2.2. Instruction modulates brain-to-brain coupling within instructor-learner dyads

Having established that social interactive learning is associated with a significant increase in brain-to-brain coupling between instructor and learner, we next sought to determine whether such coupling enhancement was modulated by Instructional Strategy and Instructional Personalization. First, results showed a main effect of Instructional Strategy in prefrontal regions (i.e., CHs 5, 6, 10, 12) at 0.45 – 0.57 Hz (*F*_s_ > 9.50, FDR corrected *p*_s_ < 0.05, *η*^2^s > 0.65). Further analyses revealed that the scaffolding strategy exhibited higher brain-to-brain coupling than the explanation strategy in all significant CHs (**Fig. 4A**). There were no effects of Instructional Strategy for other CHs and other frequency bands (*p*s > 0.05, FDR corrected). There was no significant main effect of Instructional Personalization in any CHs and at any frequency bands (*p*s > 0.05, FDR corrected).

**Figure 4.**
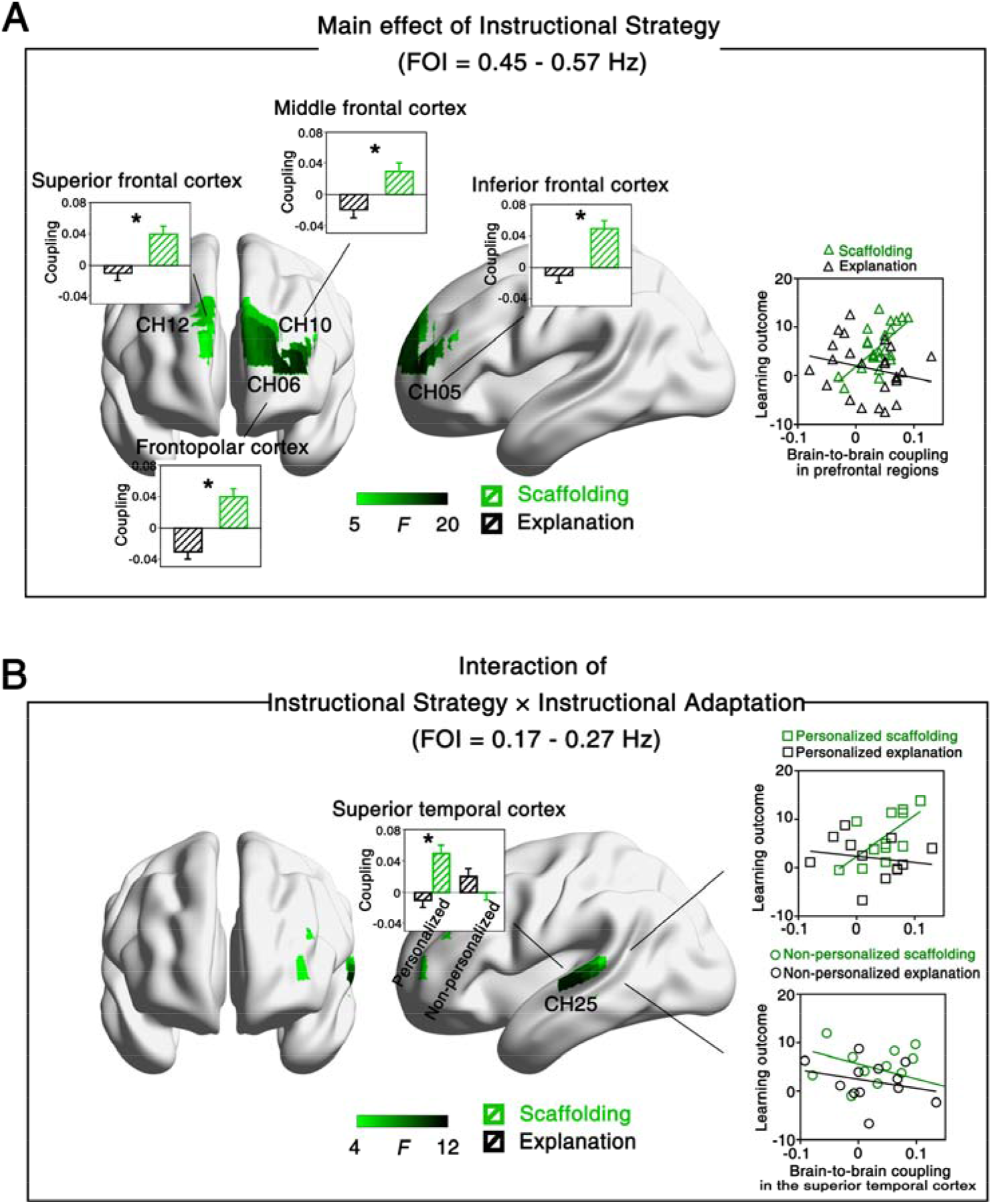
Instruction modulates brain-to-brain coupling during social interactive learning. Central: F-test maps of brain-to-brain coupling generated based on frequency-specific ANOVAs with Instructional Strategy and Instructional Personalization as independent variables. **(A)** The scaffolding condition showed higher brain-to-brain coupling in prefrontal regions than the explanation condition. Such brain-to-brain coupling predicted learning outcomes in the scaffolding condition, but not in the explanation condition (right panel). **(B)** The scaffolding condition also led to significantly larger brain-to-brain coupling in superior temporal cortex than the explanation condition, but only in the personalized instruction dyads. Brain-to-brain coupling predicted learning outcomes in the personalized scaffolding condition but not in other conditions (right panel). \**p* < 0.05. Error bars indicate standard errors of the mean.

We did, however, observe an interaction between Instructional Strategy and Instructional Personalization in the superior temporal cortex (i.e., CH 25) at 0.17 – 0.27 Hz (*F*_(1, 24)_ = 13.49, FDR corrected *p* < 0.05). Post hoc comparisons indicated that brain-to-brain coupling was significantly larger for the scaffolding condition than the explanation condition in the personalized group (*p* < 0.05), but not in the non-personalized group (*p* > 0.05, **Fig. 4B**). No significant main effects or interactions where observed in any other CHs or frequency bands of interest (*p*s > 0.05, FDR corrected).

Average brain-to-brain coupling in prefrontal regions was positively correlated with learning outcomes in the scaffolding condition (*r* = 0.65, *p* = 0.001; **Fig. 4A**, right panel) but not in the explanation condition (*r* = −0.24, *p* = 0.27), indicating that better learning was associated with stronger brain-to-brain coupling in the scaffolding condition alone. Mirroring the ANOVA results reported above, we saw that brain-to-brain coupling in superior temporal cortex only predicted learning outcomes in the personalized scaffolding condition (*r* = 0.66, *p* = 0.02; all other conditions: *r*s < −0.18, *p*_s_ > 0.27; **Fig. 4B**, right).

#### 3.2.3. Linking instructional behaviors with brain-to-brain coupling

To investigate how instructional behaviors contributed to brain-to-brain coupling, we conducted a video coding analysis for each participant dyad. Two raters independently coded videos for scaffolding behaviors vs. non-scaffolding instructional behaviors (or explanatory behaviors vs. non-explanatory instructional behaviors). For analysis, time courses of brain-to-brain coupling during the task session were first matched with video-coded instructional behaviors (**Figs. 5A–C**). Brain-to-brain coupling was then extracted for segments of each type of instructional behavior and averaged for each condition. Task-related coupling was then obtained by subtracting time-averaged brain-to-brain coupling during the rest session from the averaged coupling segments during the task session (**Figs. 5D&E**).

First, we examined whether task-related brain-to-brain coupling in prefrontal cortex detected in the scaffolding condition could be explained by scaffolding behaviors. Indeed, scaffolding behaviors induced significantly higher brain-to-brain coupling compared to the non-scaffolding instructional behaviors (*t*_(23)_ = 2.72, *p* = 0.01, Cohen’s *d* = 0.78; **Fig. 5D**, upper panel). Crucially, we also compared. However, no significant differences in brain-to-brain coupling were seen between explanatory behaviors and non-explanatory instructional behaviors in the explanation condition (*t*_(23)_ = 1.58, *p* = 0.13; **Fig. 5D**, lower panel).

Second, we compared brain-to-brain coupling for scaffolding vs. non-scaffolding instructional behaviors to test whether scaffolding behavior indeed drove the task-related brain-to-brain coupling observed in superior temporal cortex for the personalized scaffolding condition. As expected, scaffolding behaviors exhibited larger brain-to-brain coupling than non-scaffolding instructional behaviors (*t*_(11)_ = 3.19, *p* = 0.01, Cohen’s *d* = 1.18; **Fig. 5E**, upper panel). In contrast, just like in prefrontal cortex, brain-to-brain coupling did not differ between explanatory behaviors and non-explanatory behaviors in the personalized explanation condition (*t*_(11)_ = 0.91, *p* = 0.38 (**Fig. 5E**, lower panel). Moreover, there was no significant difference between instructional behaviors in either non-personalized scaffolding (**Fig. 5F**, upper panel) or non-personalized explanation conditions (**Fig. 5F**, lower panel, *t*_s_ < 1.36, *p*_s_ > 0.20).

Importantly, the effects reported here cannot be attributed to differences between conditions in terms of the mere quantity of instructional behaviors or the number of turn-takings, as evidenced by two control analyses. First, we calculated the duration ratio of instructional behaviors by quantifying the proportions of time (out of 8 minutes) when instructional behaviors occurred (Jiang et al., 2015; Pan et al., 2018). For example, if scaffolding behaviors occurred for a total of 3 minutes in an instructor-learner dyad, then the duration ratio of scaffolding behaviors should be 3/8 = 0.375. Results revealed that the duration ratio was comparable between scaffolding behaviors (0.56 ± 0.18) and non-scaffolding instructional behaviors (0.44 ± 0.18) in the scaffolding condition (*t*_(23)_ = 1.22, *p* = 0.25). Second, we compared the cumulative number of sequential turn-takings during interactive learning (for example, one turn-taking event could be that the instructor asks one question, followed by the answer from the learner). Results showed that the scaffolding strategy involved marginally more turn-takings than the explanation strategy (16.67 ± 6.54 vs. 12.08 ± 3.15; *t*_(23)_ = 2.11, *p* = 0.06). No significant correlation between the number of turn-takings and brain-to-brain coupling was detected (*r*s < 0.42, *p*s > 0.18).

In sum, brain-to-brain coupling could be explained by dynamic scaffolding behavior implemented in the instructor-learner interaction. Our complementary analyses ruled out frequency of instructional behaviors or turn-taking behavior as possible contributors to the observed brain-to-brain coupling effects.

#### 3.2.4. Decoding instructional strategy from brain-to-brain coupling

Finally, we tested the extent to which one can identify the Instructional Strategy employed by an instructor (i.e., *scaffolding* or *explanation)* based on task-related brain-to-brain coupling alone. Brain-to-brain coupling was extracted from all channel combinations that showed significantly higher brain-to-brain coupling for task vs. baseline to train the classifiers. The classifier successfully distinguished instructors who employed the *scaffolding* or *explanation* strategy with an Area Under the Curve (AUC) of 0.90, i.e., significantly exceeding chance (*p* < 0.0001, **Fig. 6A**). The decoding analysis based on task-related brain-to-brain coupling further showed that the classifier was able to distinguish instructors who employed the *scaffolding* or *explanation* strategy for the personalized condition (AUC = 0.84; *p* = 0.005, **Fig. 6B**), but not in the non-personalized condition (AUC = 0.66; *p* = 0.17, **Fig. 6C**).

**Figure 6.**
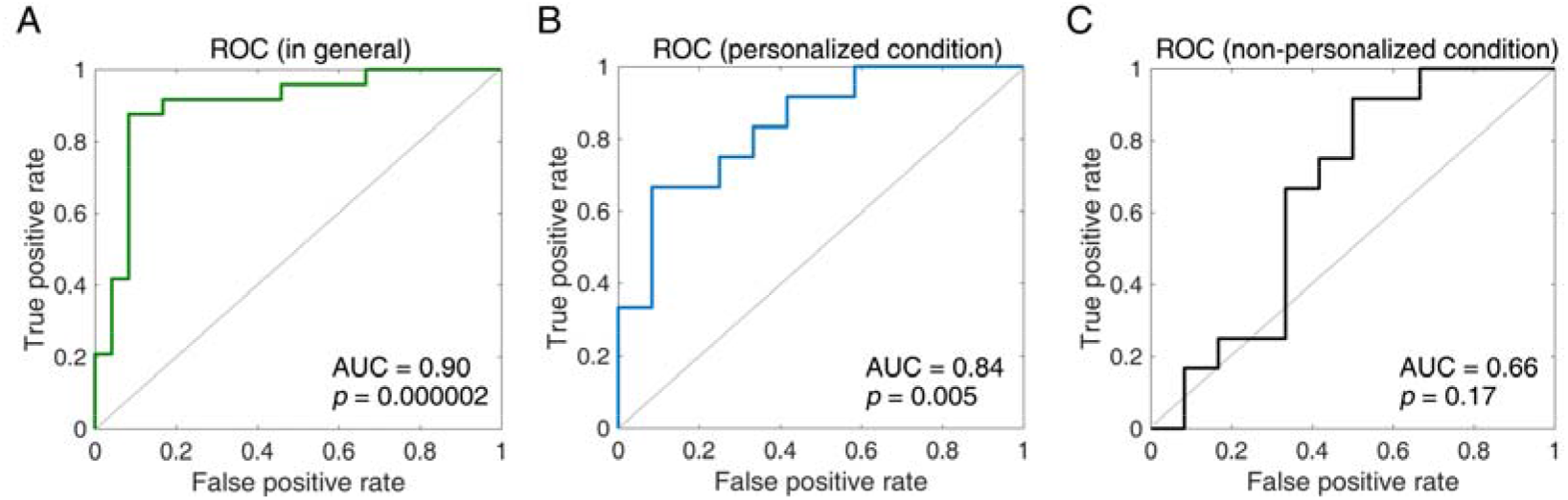
Decoding performance. The receiver operating characteristic (ROC) curve for classification distinguishing the *scaffolding* or *explanation* strategy in general **(A)**, in the personalized **(B)**, and non-personalized conditions **(C)**. Area under the curve (AUC) was calculated. Significant levels were calculated by comparing the correct AUC from the real labels with 10000 renditions of randomized labels.

Importantly, when using individual brain activation from either instructors’ or learners’ as classification features, classification performance to discriminate between the *scaffolding* and *explanation* strategies was low (AUCs < 0.66, *p*_s_ > 0.05). The decoding analysis based on the individual brain activation was also insufficient to distinguish the *scaffolding* and *explanation* strategies for both personalized (AUCs < 0.57, *p*_s_ > 0.35) and non-personalized conditions (AUCs < 0.56, *p*_s_ > 0.20).

Taken together, these results indicate that brain-to-brain coupling, as a novel yet promising neural-classification feature (Jiang et al., 2015), was suitable for decoding instructional strategy with a reasonable classification performance, particularly when the instruction was tailored to the learner (i.e., personalized vs. non-personalized). Brain-to-brain coupling further served as a better classification feature compared to individual brain activation during instructor-learner interactions.

## 4. Discussion

This study investigated how verbal instruction modulates interactive learning using an fNIRS-based hyperscanning approach, which allowed us to record brain activity from both instructors and learners *during* an instruction exchange. Twenty-four instructor-learner dyads performed a conceptual learning task in a naturalistic instruction situation where a well-trained instructor taught a learner a set of psychological concepts. We found that interactive learning induced task-related brain-to-brain coupling. Brain-to-brain coupling co-varied with learners’ subsequent learning outcomes and was significantly higher when instructors employed scaffolding tactics (e.g., asking key questions and hinting) than when they used an explanation-based teaching approach. This brain-to-brain coupling associated with scaffolding was especially prominent if instructors were informed of the learner’s knowledge level in advance. Finally, different instructional strategies could successfully be decoded based on brain-to-brain coupling alone, but, crucially, not based on individual brain activation.

Importantly, our findings were specific to the interacting instructor-learner dyads (control analysis #1) and they did not reflect the mere quantity of instructional behaviors (control analysis #2), nor the amount of turn-takings between instructor and learner (control analysis #3).

### 4.1. Using two brains to study learning and instruction

Educators have long debated which method of instruction is most conducive to learning. Several researchers have sought an answer to this question by studying learners’ neural activity during both information encoding and retrieval. However, previous studies have primarily focused on isolated individuals (e.g., Hartstra et al., 2011; Olsson and Phelps, 2007; Ruge and Wolfensteller, 2009). This poses a limitation to obtaining full insight into the learner process, especially for instruction-based learning, which relies on the dynamic instructional interaction between instructor and learner. A “second-person approach” (also termed as “hyperscanning”, i.e., measuring two brains simultaneously, Redcay and Schilbach, 2019) provides a possible way to fill this knowledge gap.

The second-person approach allowed us to quantify brain-to-brain coupling between the instructor and the learner, and possibly capture the continuous, meaningful alignment of interpersonal neural processes. It has been proposed that such neural alignment facilitates the matching of the temporal structure of inputs and optimizes the learning process (Leong et al., 2017). Our findings suggest that brain-to-brain synchrony is pedagogically relevant. First, brain-to-brain coupling was correlated with learning outcomes, strongly indicating its functional significance. Second, brain-to-brain coupling was successfully used to decode instructional approaches with a good classification performance.

To our knowledge, we are the first to use activity from two brains as opposed to one to decode instructional strategies. We found that brain-to-brain coupling served as a better neural-classification feature in contrast with individual brain activity. This finding was in line with recent advances; for example, a recent study found that brain-to-brain coupling yielded higher predictive power for learning outcomes compared to single-brain measures (Davidesco et al., 2019). A possible explanation for this is that non-neuronal artifacts are systematic in individual brain activity (Zhang et al., 2016), while such artifacts are not consistent across brains. Indeed, brain-to-brain coupling has been reported to have higher signal-to-noise than single-brain measures (Parkinson et al., 2018). Moreover, measuring coupling across brains can provide complementary information that cannot be revealed by conventional individual brain measures (Balconi et al., 2017; Simony et al., 2016). Compared to single-brain activity, brain-to-brain coupling could be more sensitive when tracking ongoing social interactions because it considers the neural dynamics from all interacting agents simultaneously. In sum, there are several benefits of recording activity from two brains (versus one brain) to study learning and instruction.

### 4.2. The role of prefrontal and temporal cortices in brain-to-brain coupling

The modulatory effects of instruction on brain-to-brain coupling were concentrated in prefrontal and superior temporal cortices. This is in line with prior fNIRS-based hyperscanning studies that found that brain-to-brain coupling in prefrontal cortices (PFC; Holper et al., 2013; Pan et al., 2018; Takeuchi et al., 2017) and temporoparietal regions (Zheng et al., 2018) predicted learning outcomes following instruction. PFC has been associated with a wide range of human cognitive functions. Specific to hyperscanning, PFC has been implicated in cooperation (Cheng et al., 2015), competition (Liu et al., 2015), and emotion regulation (Reindl et al., 2018). In this study, the scaffolding process might require constant collaborative interaction between instructor and learner, a process for which prefrontal areas are heavily recruited.

Superior temporal cortex (STC), like PFC, has been associated with many cognitive functions that are relevant for learning, and social cognition more broadly. For example, STC is a key area for theory of mind or mentalizing (Baker et al., 2016), and has been implicated in social perception and action observation (Thompson and Parasuraman, 2012). While the exact role of STC in brain-to-brain coupling during learning cannot be inferred based on the present findings, it is possible that brain-to-brain coupling in this area reflects the shared intentionality or mental state between instructor and learner, or a process whereby instructors need to infer the understanding of the learner such that instruction can be adapted or personalized accordingly (Zheng et al., 2018).

Another possibility is that the correlation between brain-to-brain synchrony and learning outcomes in STC and PFC can be accounted for in terms of the ability of the instructor and learner to predict each other’s mental states and utterances throughout the interaction. Prior fMRI studies investigating speaker-listener brain-to-brain coupling found that brain activity was more correlated between speakers and listeners in STC for more predictable speech (Dikker et al., 2014) and PFC brain-to-brain coupling has been associated with information retention (Stephens et al., 2010). Both PFC and STC have been found crucial for temporal predictive encoding and integration of behavior (Amoruso et al., 2018; Yang et al., 2015) and recent models attribute a large role to predictive coding in explaining interpersonal alignment at both the neural and the behavioral level (Garrod and Pickering, 2010; Shamay-Tsoory et al., 2019).

### 4.3. Linking brain imaging findings to pedagogical practice

As the Chinese educator Confucius suggested, appropriate instruction matters during instructor-learner interactive learning (Chen, 2007). Several theoretical models have been proposed aiming at improving pedagogy. These models include explanation-based and constructivism-based theories, both of which have been shown demonstrated to support learning (Chi, 2013).

As laid out in the introduction, an explanation-based approach puts emphasis on information clarification and aims at providing prefabricated explanatory information to the learner. Explanation is a conventional strategy used in classroom instruction (Leinhardt and Steele, 2005), human tutoring (Chi et al., 2004), cooperative learning (Webb et al., 2006), and skill acquisition (Renkl et al., 2007). In a constructivism-based approach, in contrast, the instructor is encouraged to provide support (i.e., scaffolding) tailored to the needs of the learner (Kleickmann et al., 2016). In this framework, instructional modulation of learning arises from exogenous constructivist instruction (Jumaat and Tasir, 2016). Arguably, our findings favor a constructivism-based model: brain-to-brain coupling during interactive learning was primarily driven by the moments of scaffolding behaviors, a central feature of a constructivist approach to instruction-based learning. It is important to note that our results do not warrant the conclusion that explanation-based instruction is not useful: This would go against decades of research showing that people do learn from explanations (Chi et al., 2004; Leinhardt and Steele, 2005; Renkl et al., 2007; Webb et al., 2006).

Our findings can also be interpreted within the context of the *Interactive-Constructive-Active-Passive* (ICAP, Chi and Wylie, 2014) framework. The ICAP framework defines a set of cognitive engagement activities, which can be categorized into *Interactive, Constructive, Active,* and *Passive* modes, based on learners’ behaviors. The four modes correspond to different cognitive processes (Lam and Muldner, 2017): *Interactive* engagement corresponds to the cognitive process of co-creating knowledge (e.g., dialogues); *Constructive* engagement involves creating knowledge (e.g., explaining in one’s own words); *Active* engagement involves emphasizing or selecting knowledge (e.g., copying notes); *Passive* engagement involves storing knowledge (e.g., watching and listening to the instructor). The ICAP hypothesis proposes that the learning increase from *Passive* to *Active* to *Constructive* to *Interactive.* In the current study, although both strategies involved interactive engagement, the scaffolding strategy could additionally invoke constructive engagement whereas the explanation strategy could invoke relatively passive engagement in the learners (as summarized in **Fig. 7**). Consistent with the ICAP, learning outcomes were better in the scaffolding than the explanation strategies, i.e., (*Interactive* + *Constructive*) > (*Interactive* + Passive). What’s more, one can argue that our results extend the theoretical framework of ICAP by showing that the four components proposed should not be treated in isolation: real-life instruction is a complex activity and generally engages several cognitive components. Our findings suggest that instructors should consider including and combining more interactive and constructive engagements.

**Figure 7.**
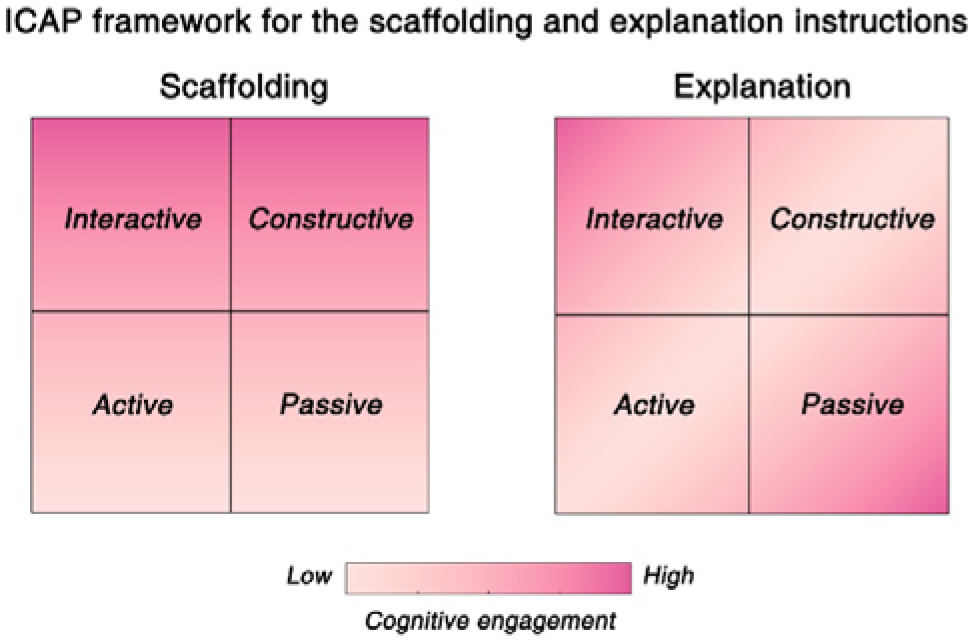
Interactive-Constructive-Active-Passive (ICAP) framework for the scaffolding and explanation instructions. The scaffolding instruction elicits more interactive and constructive responses, whereas the explanation instruction elicits more interactive and passive responses.

### 4.4. Conclusions

Recording brain activity from multiple participants simultaneously in ecologically valid settings is a nascent but promising field of research. We investigated interactive learning using fNIRS hyperscanning in a naturalistic learning situation, and found that verbal instruction modulates learning via brain-to-brain coupling between instructors and learners, which was driven by dynamic scaffolding representations. Importantly, brain-to-brain coupling was effective to discriminate between different instructional approaches and predict learning outcomes. Together, our findings suggest that brain-to-brain coupling may be a pedagogically informative implicit measure that tracks learning throughout ongoing dynamic instructor-learner interactions.

## Contribution

Y. P., C. Y., and Y. H. designed the experiment. Y. P., Y. Z., and C. Y. performed the study. Y. P. analyzed the data. Y. P., S. D., P. G., Y. Z., C. Y., and Y. H. wrote the manuscript.

## Competing financial interests

The authors declare no competing financial interests.

## Funding

This work was sponsored by the National Natural Science Foundation of China (31872783), the General Project of Humanities and Social Sciences of the Ministry of Education (19A10332020), the China Scholarship Council (201706140082), and the outstanding doctoral dissertation cultivation plan of action of East China Normal University (YB2016011).

## Notes

#### Summary of Updates

Title and abstract updated.

